# MicrobiomeGWAS: a tool for identifying host genetic variants associated with microbiome composition

**DOI:** 10.1101/031187

**Authors:** Xing Hua, Lei Song, Guoqin Yu, James J. Goedert, Christian C. Abnet, Maria Teresa Landi, Jianxin Shi

## Abstract

The microbiome is the collection of all microbial genes and can be investigated by sequencing highly variable regions of 16S ribosomal RNA (rRNA) genes. Evidence suggests that environmental factors and host genetics may interact to impact human microbiome composition. Identifying host genetic variants associated with human microbiome composition not only provides clues for characterizing microbiome variation but also helps to elucidate biological mechanisms of genetic associations, prioritize genetic variants, and improve genetic risk prediction. Since a microbiota functions as a community, it is best characterized by beta diversity, that is, a pairwise distance matrix. We develop a statistical framework and a computationally efficient software package, microbiomeGWAS, for identifying host genetic variants associated with microbiome beta diversity with or without interacting with an environmental factor. We show that score statistics have positive skewness and kurtosis due to the dependent nature of the pairwise data, which makes P-value approximations based on asymptotic distributions unacceptably liberal. By correcting for skewness and kurtosis, we develop accurate *P*-value approximations, whose accuracy was verified by extensive simulations. We exemplify our methods by analyzing a set of 147 genotyped subjects with 16S rRNA microbiome profiles from non-malignant lung tissues. Correcting for skewness and kurtosis eliminated the dramatic deviation in the quantile-quantile plots. We provided preliminary evidence that six established lung cancer risk SNPs were collectively associated with microbiome composition for both unweighted (P=0.0032) and weighted (P=0.011) UniFrac distance matrices. In summary, our methods will facilitate analyzing large-scale genome-wide association studies of the human microbiome.

## Introduction

The human body is colonized by bacteria, viruses and other microbes that exceed the number of human cells by at least 10-fold and that exceed the number of human genes by at least 100-fold. The relationship between a person and his or her microbial population, termed the microbiota, is generally mutualistic. The microbiota may promote human health by inhibiting infection by pathogens, conditioning the immune system, synthesizing and digesting nutrients, and maintaining overall homeostasis. The microbiome, which is the collection of all microbial genes, can be investigated through massively parallel, next-generation DNA sequencing technologies. By amplifying and sequencing highly variable regions of 16S ribosomal RNA genes that are present in all eubacteria, cost-effective and informative microbiome profiles down to the genus level are obtained.

The human microbiome has been associated with diseases, including obesity^1^, inflammatory bowel disease (IBD)^2^, colorectal cancer^3^ and breast cancer^4^. Thus, identifying factors that have a sustained impact on the microbiome is fundamental for elucidating its role in health conditions and for developing treatment strategies. Increasing evidence suggests that microbiome composition at a specific site of the human body is impacted by environmental factors^5,6^, host genetics^7,8^, and possibly by their interactions. In the mouse, quantitative trait loci (QTL) studies have identified loci contributing to the variation of the gut microbiome using linkage analysis^9,10^. Recently, Goodrich et al.^11^ systematically investigated the heritability of the human gut microbiome by comparing monozygotic twins to dizygotic twins and found substantial heritability in different microbiome metrics, suggesting the important role of host genetics on gut microbiome diversity. Associations between individual host genetic variants and microbiome taxa abundances have also begun to emerge in other human samples^7,8,12^. These studies suggest that genome-wide association studies (GWAS) have great potential to identify host genetic variants associated with microbiome diversity.

GWAS of complex human diseases have identified many risk SNPs; however, the biological mechanisms are largely unknown for the majority of the risk SNPs. QTL studies of intermediate traits, e.g., gene expression^13,14^, DNA methylation^15,16^, chromatin structure^17,18^, and metabolite production^19,20^, have provided useful insights on biological mechanisms of the GWAS findings. The human microbiome at a specific body site is another important and informative intermediate trait for interpreting GWAS signals. Knights et al.^8^ reported that a risk SNP for IBD located in *NOD2* was associated with the relative abundance of *Enterobacteriaceae* in the human gut microbiome. Tong et al.^7^ show that a loss-of-function allele in *FUT2* that increases the risk of developing Crohn’s Disease (CD) may modulate energy metabolism of the gut microbiome. In both examples, the microbiome is a potential intermediate for explaining the association between risk SNPs and disease risks, although a formal mediation analysis is required based on samples with genotype, microbiome, and disease status data. Moreover, identifying microbiome-associated host genetic variants has the potential to prioritize SNPs for discovery and to improve the performance of polygenetic risk prediction.

Three types of microbiome metrics can be derived as phenotypes for GWAS analysis. First, for each taxon at a specified taxonomic level (phylum, class, order, family, genus, and species), we calculate the relative abundance (RA) of the taxon as the ratio of the number of sequencing reads assigned to the taxon to the total number of sequencing reads. In 16S ribosomal RNA sequence profiles, approximately 100-200 taxa with average RAs ≥0.1% (from the phylum level to the genus level) across samples are abundant enough for QTL analysis. One can perform a Poisson regression to examine the association between RA of each taxon and each SNP. Significant associations are identified using Bonferroni correction (*P*<5×10^−8^/200=2.5×10^−10^) or by controlling FDR at an appropriate level. Second, multiple alpha-diversity metrics^21^ can be calculated to reflect the richness (e.g., number of unique taxa) and evenness of each microbiome community after a procedure called rarefication, that eliminates the dependence between estimated alpha diversity and the variable total number of sequencing reads across subjects. Once alpha-diversity metrics are derived, one may perform standard GWAS with alpha diversity as the phenotype using linear regression.

Because a microbiota functions as a community, the most important analysis for a microbiome GWAS may be by assessing the complete structure of the community by using a pairwise microbiome distance matrix (or beta-diversity) of the microbial community. Microbiome distances can be defined in different ways, based on using phylogenetic tree information or each taxon’s abundance information. Bray–Curtis dissimilarity^22^ quantifies the difference between two microbiome communities using the abundance information of specific taxa. UniFrac^23-25^ is another widely used distance metric. Unlike the Bray–Curtis dissimilarity metric, UniFrac compares microbiome communities by using information on the relative relatedness of each taxon, specifically by phylogenetic distance (branch lengths on a phylogenetic tree). UniFrac has two variants: the weighted UniFrac^24^ that accounts for the taxa abundance information, and the unweighted UniFrac^23^ that only models the information of presence or absence. Recently, a generalized UniFrac distance metric^26^ was developed to automatically appreciate the advantages of weighted and unweighted UniFrac metrics and was shown to provide better statistical power to detect associations between human health conditions and microbiome communities. GWAS based on a microbiome distance matrix aims to identify host SNPs associated with microbiome composition. Intuitively, the microbiome distances tend to be smaller for pairs of subjects with similar genotypic values at the associated SNP. In addition, it is also of great interest to identify host SNPs that interact with an environment factor to affect microbiome composition. Importantly, beta diversity is temporally more stable compared with RA of taxa and alpha-diversity metrics based on the data from the Human Microbiome Project^27^ (data not shown), suggesting smaller power loss for a GWAS due to temporal variability. To our knowledge, no statistical methods or software packages have been designed to efficiently analyze microbiome GWAS data using distance matrices as phenotypes.

In this paper, we develop a statistical framework and a computationally efficient package, microbiomeGWAS, for analyzing microbiome GWAS data. Our package allows the detection of host SNPs with a main effect or interaction with an environment factor, i.e. host SNPs interacting with an environment factor to affect the microbiome composition. We calculate the variance of the score statistics by appropriately considering the dependence of the pairwise distances. Importantly, we show that the score statistics have positive skewness and kurtosis due to the dependence in pairwise distances, which makes the approximation of small *P*-values based on the asymptotic distribution too liberal, which easily yields false positive associations. Resampling methods, e.g. bootstrap or permutation, are computationally prohibitive for accurately approximating small P-values. We propose to improve the tail probability approximation by correcting for skewness and kurtosis of the score statistics. Numerical investigations demonstrate that our method provides a very accurate approximation even for *P*=10^−7^. MicrobiomeGWAS runs very efficiently, taking 36 minutes for analyzing main effects and 69 minutes for analyzing both main and interaction effects for a study with 2000 subjects and 500,000 SNPs using a single core. MicrobiomeGWAS can be freely downloaded at https://github.com/lsncibb/microbiomeGWAS.

We illustrate our methods by applying microbiomeGWAS to non-malignant lung tissue samples (*N* = 147) in the Environment And Genetics in Lung cancer Etiology (EAGLE) study^28,29^. Because smoking may alter microbiome composition, we tested both main effect and gene-smoking interaction effect. When P-values were calculated based on asymptotic distributions, the quantile-quantile (QQ) plots strongly deviated from the uniform distribution. Also, nine loci achieved genome-wide significance based on asymptotic approximations. Correcting for skewness and kurtosis eliminated the inflation and also the genome-wide significance of these loci. However, we provide evidence that the established lung cancer risk SNPs are associated with lung microbiome composition.

## Material and Methods

### A score statistic for testing main effect

Suppose that we have a set of *N* subjects genotyped with SNP arrays. For notational simplicity, we consider only one SNP with minor allele frequency (MAF) denoted as *f*. Our interest centers on testing whether the genotype of the SNP is associated with microbiome composition. Let *g_n_* = 0,1,2 represent the number of the minor alleles for the *n^th^* subject. We assume that the 16S rRNA gene of microbiota from a target site (e.g., gut) has been sequenced for these samples. Let *d_ij_* be the microbiome distance between the *i^th^* and *j^th^* subjects and ***D*** be the distance matrix.

Intuitively, if the SNP is associated with the microbiome composition, the microbiome distances tend to be smaller for subject pairs with similar genotypic values, as is illustrated in Figure 1. For *N* subjects, *N*(*N* − l)/2 pairs can be divided to three groups with genetic distance 0, 1 and 2. For example, a pair of subjects with genotype (AA, AA) or (BB, BB) has genetic distance 0; a pair of subjects with genotype (AA, BB) or (BB, AA) has genetic distance 2; all other pairs have genetic distance 1. Apparently, we expect the microbiome distance to be positively correlated with genetic distance for subject pairs. We define *G_ij_= |g_i_ − g_j_*| as the genetic distance for a pair of subjects (*i,j*). We assume *d_ij_* = *α* + *β_M_G_ij_ + ε_ij_* for all pairs of subjects. The score statistic for testing *H*_0_*: β_M_* = 0 (main effect) vs. *β_M_* > 0 is derived by maximizing Σ*_i<j_*(*d_ij_* − *α* − *β_M_G_ij_*)^2^:

**Figure 1.**
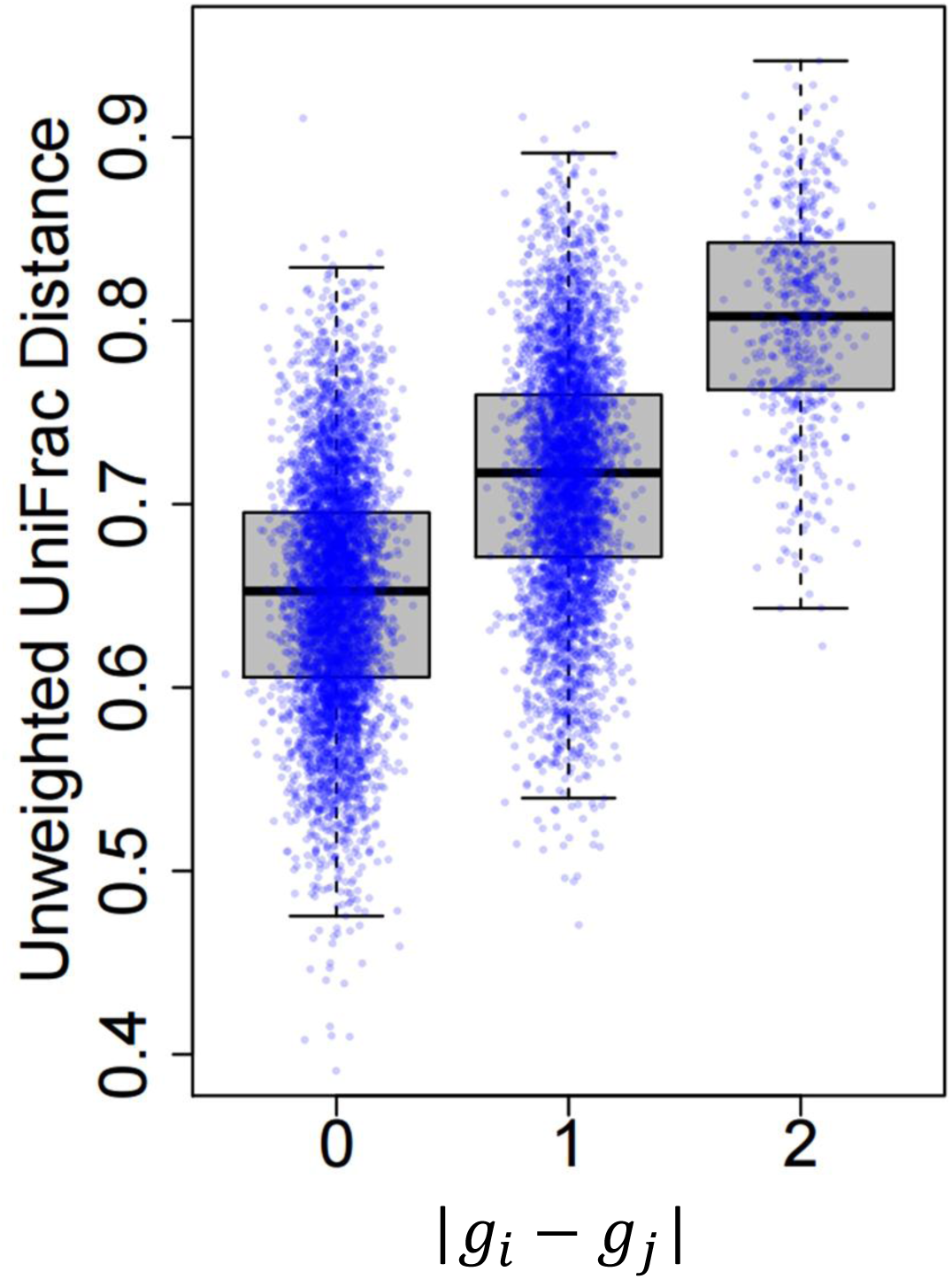
Microbiome distances are positively correlated with genetic distances at an associated SNP.

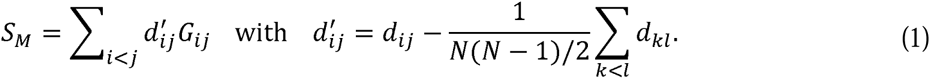

The variance *Var*_0_(*S_M_*|**D**) under *H*_0_*:β_M_* = 0 is calculated by considering the dependence in (*G_ij_*, *G_kl_*) and conditioning on the distance matrix ***D***. Briefly, we have 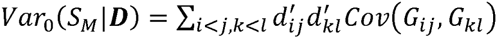. When (*i, j, k, l*) are distinct, *G_ij_* and *G_kl_* are independent, i.e. *Cov*(*G_ij_*, *G_ij_*) = 0. Some algebra leads to

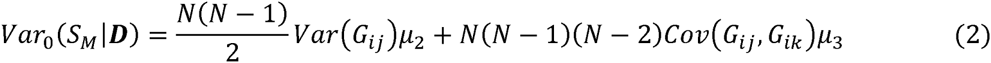

where

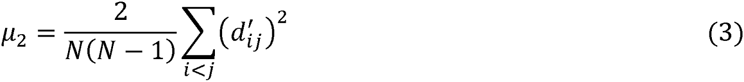

and

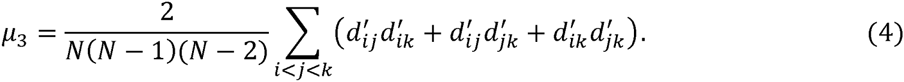

The details for calculating *Var*(*G_ij_*) and *Cov*(*G_ij_, G_ik_*) are in **Appendix A**. The variance-normalized score statistic 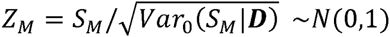 under *H*_0_ asymptotically.

In analyses of real data, we typically have to adjust for covariates, including demographic variables and principal component analysis (PCA) scores derived based on genotypes to eliminate potential population stratification. Let *X_i_* = (*x_i_*_1_, …, *x_iv_*) denote the *v* covariates for the *i^th^* subject. We assume 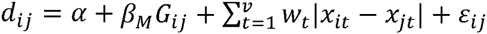. Define 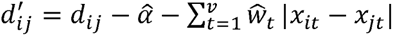 with 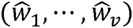 being estimated under *H_0_: β_M_* = 0. It is straightforward to verify that the score equation for *β_M_* evaluated at *H_0_: β_M_* = 0 is 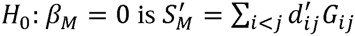. We can similarly derive the variance 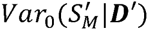 and the normalized score statistic 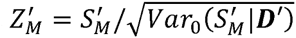. Here, ***D′*** denotes the residue distance matrix with 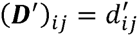.

### A score statistic for testing gene-environment interaction

Let *E_i_* denote an environmental variable. Define Δ*_ij_*·= *|g_i_E_i_ − g_j_E_j_*|. We extend the statistical framework to detect the SNP-environment interaction by assuming *d_ij_* = α + *β_M_ G_ij_*+*β_E_*|*E_i_* − *E_j_*| + *β_I_*Δ*_ij_* + *ε_ij,_* where *β_M_* denotes the main genetic effect, *β_I_* denote the additive gene-environment effect and *β_E_* denotes the main effect of the environmental factor. We consider testing the null hypothesis that the SNP is not associated with microbiome composition either directly or by interacting with *i. e. H*_0_*: β_M_* = *β_I_* = 0. The alternative hypothesis is *H*_1_*: β_M_* > 0 or *β_I_* > 0.

Again, we estimate *β_M_* and *α* under *H*_0_ and calculate 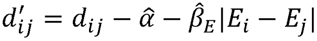. The score equations evaluated under *H*_0_ are 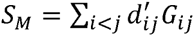 for *β_M_* and 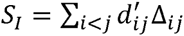 for *β_I_*. Similar to (2), we derive the variance *Var*_0_(*S_I_\****D***′) by accounting for the dependence in (Δ*_ij_*, Δ*_kl_*):

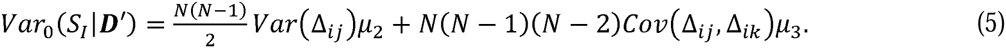

Let 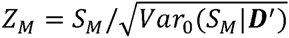 and 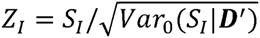. Asymptotically, *Z_M_*~*N*(0,1) and *Z_I_*~*N*(0,1) under *H*_0_.

In **Appendix B**, we derive

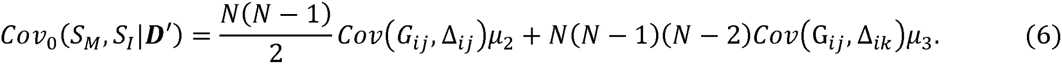

The correlation *ρ = Cor*_0_(*Z_M_*,*Z_I_\****D***′) is calculated as 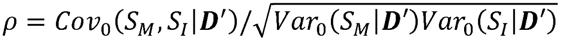. Asymptotically, (*Z_M_, Z_I_*) follows a bivariate normal distribution with a correlation matrix 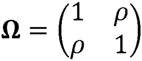 In **Appendix C**, we derive a statistic for jointly testing *H_0_*: *β_M_ = β_I_* = 0 vs.: *H*_1_ > 0 or *β_I_* > 0. Briefly, the 2D plane is partitioned to four parts (**Figure 2**). The joint statistic is derived as

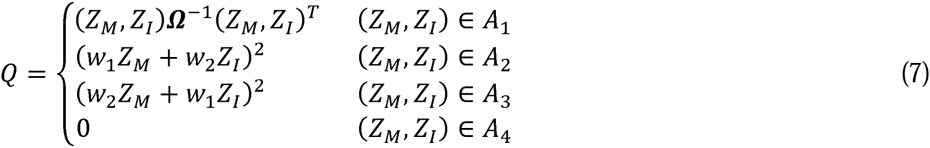

where *w*_1_ = (*θ −* 1*/θ*)/2, *w*_2_ = (*θ +* 1*/θ*)/2 and 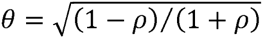. The asymptotic P-value is calculated as

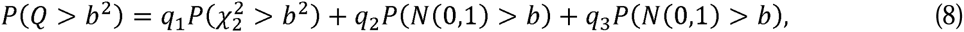

where 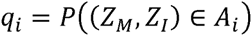.

**Figure 2.**
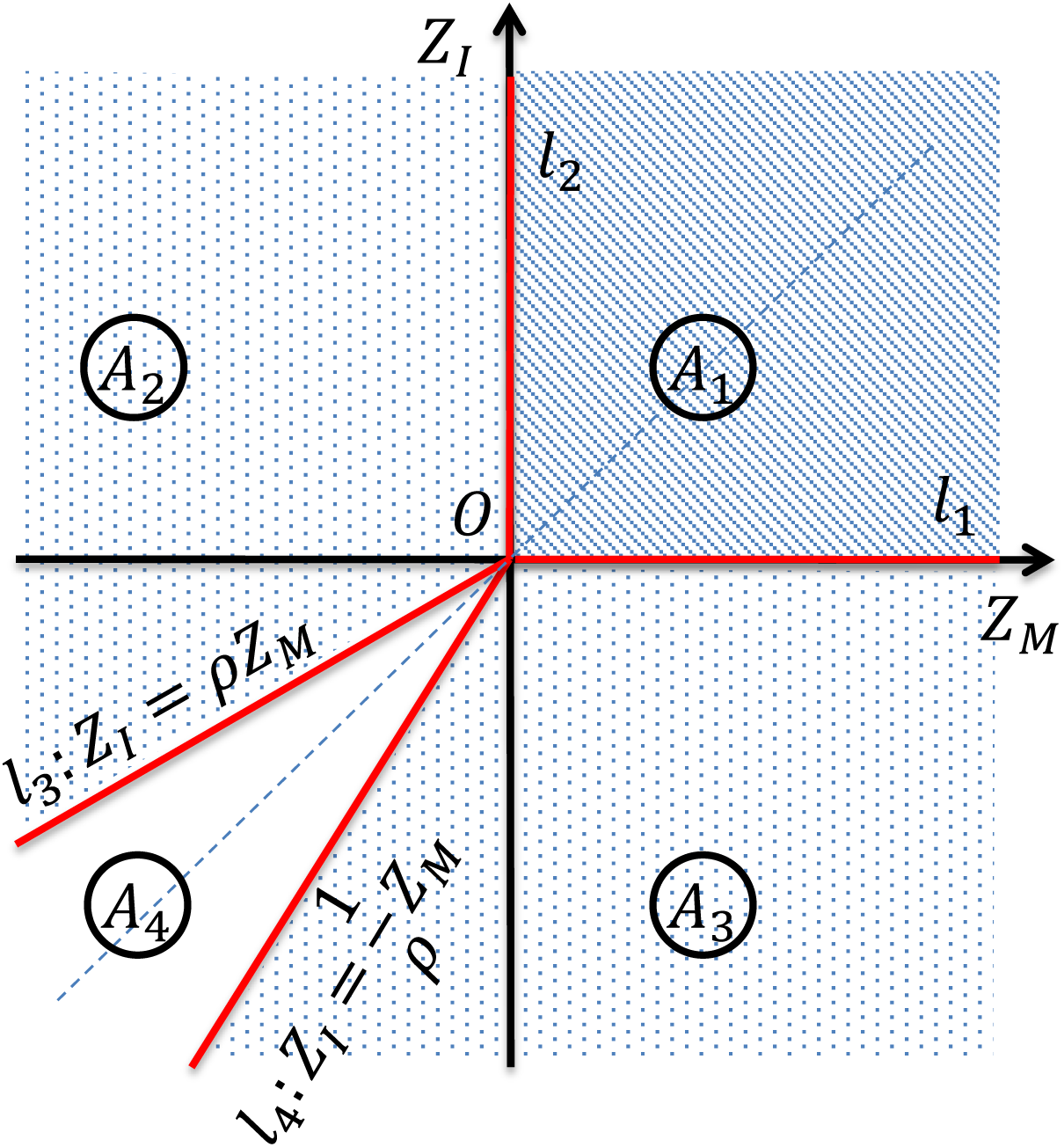
Define the joint test for testing *H*_0_: *β_M_ = β_I_* = 0 vs. *β_M_* > 0 *or β_I_* > 0. We assume that *Z_M_* ~ *N*(0,1), *Z_I_* ~ *N*(0,1) and *Cor*(*Z_M_*, *Z_I_*) = *ρ* under *H*_0_. Details are in **Appendix C**.

### Improved *P*-value approximations by correcting for skewness and kurtosis

Theoretic investigation suggests that the score statistics *Z_M_* and *Z_I_* have a positive skewness, which makes the tail probability approximations based on the asymptotic distribution *N*(0,1) unacceptably liberal (**Figures 3A and 3B**). In a numeric example with skewness *γ* = 0.2, *P*(*Z* > 5) =2.9×10^−7^ based on *N*(0,1), which is approximately two orders of magnitude more significant than P=3.9×10^−5^ based on 10^8^ permutations. The significance inflation becomes worse for smaller P-values and larger skewness *γ.* Similar but more tedious calculations suggest that both statistics have positive kurtosis, making the approximation based on *N*(0,1) even worse. One possible solution is to approximate tail probabilities using permutations or bootstrap. However, these resampling methods are computationally prohibitive for testing millions of common SNPs in a large-scale study.

**Figure 3.**
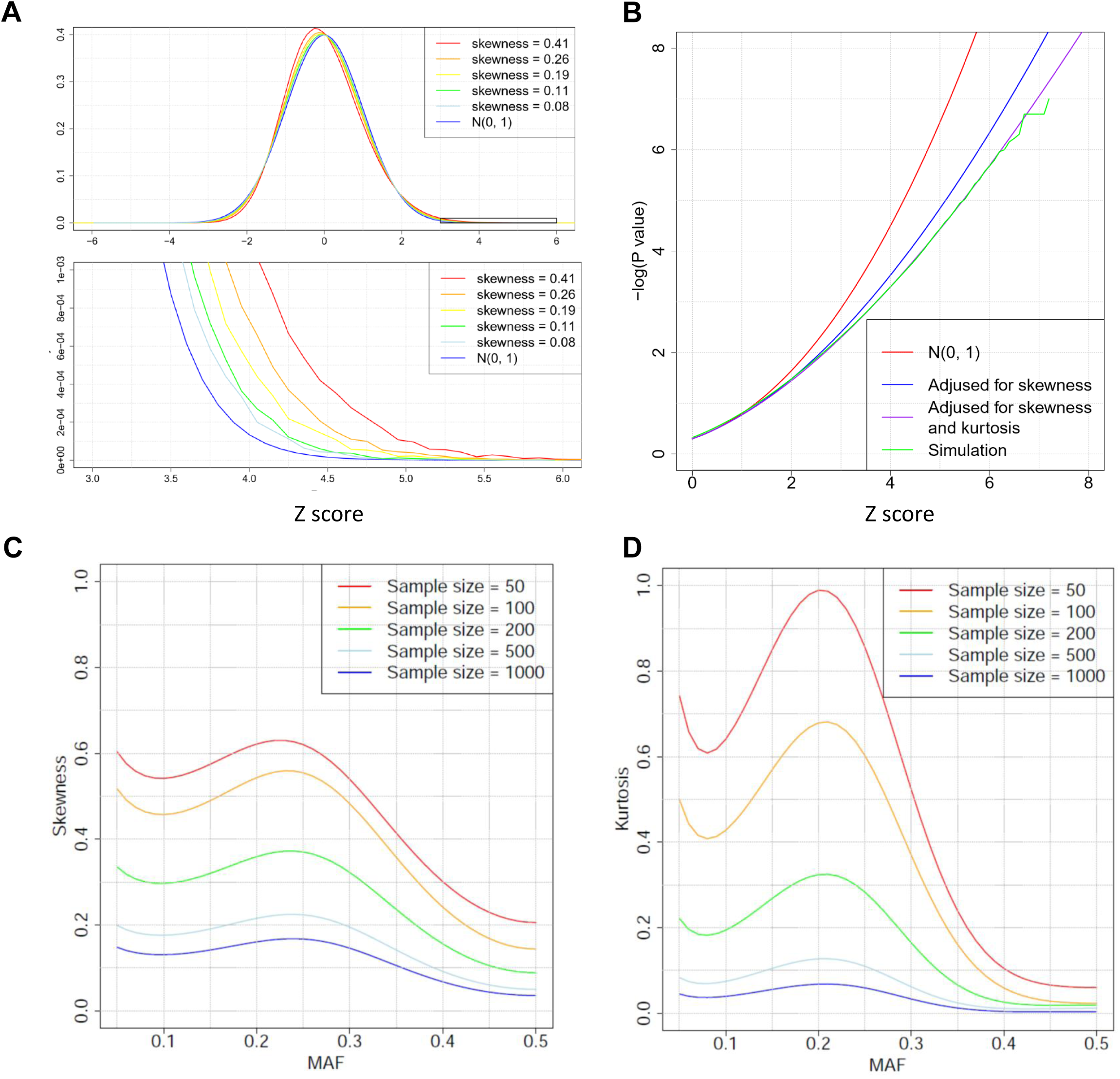
Correcting tail probabilities for skewness and kurtosis. (A) The standard normal distribution *N*(0,1) and an approximately normal distribution with positive skewness. The skewness has big impact when calculating the tail probability *P*(*Z > b*) for a large value of *b.* (B) Numerical evaluation of tail probability approximation for *Z_M_.* We used the unweighted UniFrac distance matrix of 500 samples from the American Gut Project (AGP). For each value of *b*(*>* 0), we calculated P-values *P*(*Z_M_ > b*) based on *N*(0,1), skewness correction, both skewness and kurtosis correction, and 10^8^ simulations. (C) Skewness depends on MAF of SNPs and the sample size of the study, calculated based on the weighted UniFrac distance matrix in AGP data. (D) Kurtosis depends on MAF of SNPs and the sample size, calculated based on the weighted UniFrac distance matrix in AGP data.

To address this problem, we calculated the skewness and kurtosis of the score statistics under *H*_0_ (**Appendix D**). We propose to improve the tail probability approximation *P*_0_(*Z > b*) by correcting for the skewness and kurtosis, following the skewness correction in linkage analysis^30,31^. Technical details are provided in **Appendix E**. Correcting for both skewness and kurtosis leads to an approximation

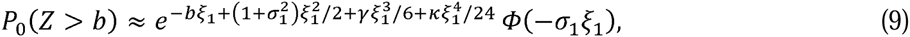

where *ξ*_1_ satisfies 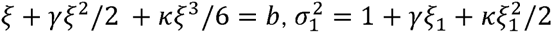, and *Φ*(·) is the cumulative distribution function of *N*(0,1). Correcting for skewness but ignoring kurtosis (i.e., assuming *κ* = 0) leads to an approximation

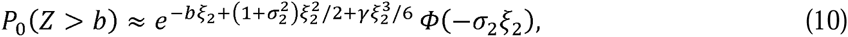

where 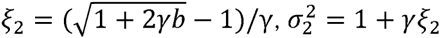. Numerical results presented in **Figure 3B** demonstrate that (9) works very well.

Given the distance matrix ***D****, γ_Μ_* ∝ 1/*N*^1/2^, *γ_I_* ∝ 1/*N*^1/2^, *k_m_* ∝ 1/*N* and *κ_I_* ∝ 1/*N* (**Appendix D**). Thus, skewness decays much more slowly with sample size *N* than kurtosis (**Figures 3C and 3D**). Thus, even for a large study with thousands of samples, correcting for skewness is necessary for accurately evaluating tail probabilities. Importantly, both skewness and kurtosis highly depend on the MAF, suggesting that the impact of skewness and kurtosis is different across SNPs with different MAF. Numerical studies (**Figures 3C and 3D**) show that skewness and kurtosis are minimized when MAF=0.5 and maximized when MAF≈0.2-0.3.

Finally, we discuss how to approximate the tail probability of *Q* in (7) for testing *H_0_: β_M_ = β_I_* = 0 by correcting for non-normality in *Z_M_* and *Z_I_*. When (*Z_M_*, *Z_I_*) ∈ *A*_2_ (or *A*_3_), we calculate the skewness *E*(w_1_*Z_M_* + w_2_*Z_M_*)^3^ and the kurtosis *E*(w_1_*Z_M_* + w_2_*Z_I_*)^4^ − 3 and use (9) to approximate *P*(w_1_*Z_M_* + w_2_*Z_I_*>*b*). When (*Z_M_,Z_I_*) ∈ *A*_1_, we first approximate their marginal *P*-values as *p_M_* and *p_I_* by (9), then calculate the normal quantile *z_M_* = *Φ*(1 − *p_M_*) and *z_I_* = *Φ*(1 − *p_I_*). Because the correction primarily impacts the tails of the distributions, the correlation between the two statistics will remain roughly unchanged, i.e., *cor*_0_(*Z_M_,Z_I_*) ≈ *cor*_0_(*z_M_, Z_I_*). Thus, when (*Z_M_,*) ∈ *A*_1_ the tail probability is approximated as 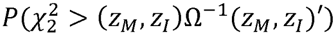.

## Results

### Simulation results

The main purpose of simulations was to investigate the type-I error of *Z_M_* (for testing main genetic effect), *Z_I_*(for detecting SNP-environment interactions) and *Q* (for detecting either main genetic effect or SNP-environment effect or both). Simulations were performed under different combinations of sample size, MAF and microbiome distance matrices. To make simulations realistic, we used an unweighted distance matrix of the fecal microbiome samples with the 16S rRNA V4 region sequences from the American Gut Project (AGP). The OTU table rarefied to 10,000 sequence reads per sample, as well as metadata, was downloaded from the AGP website. Samples with less than 10,000 sequence reads were excluded from analysis. The weighted and the unweighted UniFrac distance matrices were generated in the Quantitative Insights Into Microbial Ecology^21^ (QIIME) pipeline. Because antibiotics may substantially change microbiome composition to generate outliers that may distort the null distribution, we excluded samples with self-reported history of antibiotic usage within one month. After quality control, 1879 subjects remained for analysis. In simulations, we randomly selected *N* samples for a given sample size *N.*

For each setting, the type-I error rates were evaluated based on 10^8^ simulations under *H*_0_. For the interaction test and the joint test, the binary environment factor had a frequency of 50% and was simulated independent of the SNP. The type-I error rates are summarized in Table 1 for weighted UniFrac distance matrix. The skewness and kurtosis are reported in Figures 3C and 3D. The statistics adjusted for skewness and kurtosis have accurate type-I error rates while the statistics without adjustment have unacceptably high type-I error rates. As sample size increases, the impact of skewness and kurtosis decreases. However, even for a study with *N* = 1000, the type-I error rates are still seriously inflated. The results for the unweighted UniFrac distance matrix and for MAF=0.5 are reported in **Table S1**.

**Table 1:** Type-I error rates estimated based on 10^8^ simulations. Minor allele frequency = 20%. Simulations were based on the weighted UniFrac distance matrix of the gut microbiome data from the American Gut Project. Reported are the type-I error inflation factor. A value greater than 1 indicates an inflated type-I error.

| | N | $Z_M$ | | | $Z_I$ | | | $Q$ | | |
| --- | --- | --- | --- | --- | --- | --- | --- | --- | --- | --- |
| | | $\alpha = 10^{-3}$ | $10^{-5}$ | $10^{-7}$ | $10^{-3}$ | $10^{-5}$ | $10^{-7}$ | $10^{-3}$ | $10^{-5}$ | $10^{-7}$ |
| asymptotic approximation | 100 | 5.5 | 51.6 | 610.0 | 4.7 | 36.1 | 342.8 | 7.3 | 80.9 | 1148.0 |
|  | 200 | 3.7 | 23.0 | 187.3 | 3.1 | 15.8 | 105.5 | 4.6 | 33.0 | 316.7 |
|  | 500 | 2.4 | 9.4 | 45.2 | 2.1 | 6.7 | 25.5 | 2.8 | 11.9 | 64.1 |
|  | 1000 | 2.0 | 5.7 | 21.3 | 1.8 | 4.4 | 14.0 | 2.2 | 6.9 | 28.5 |
| adjusted for skewness and kurtosis | 100 | 1.0 | 1.2 | 0.7 | 1.0 | 1.1 | 0.6 | 1.0 | 1.5 | 2.0 |
|  | 200 | 1.0 | 1.1 | 1.0 | 1.0 | 1.1 | 0.7 | 0.9 | 1.3 | 1.8 |
|  | 500 | 1.0 | 1.1 | 1.3 | 1.0 | 1.0 | 0.9 | 0.9 | 1.0 | 1.7 |
|  | 1000 | 1.0 | 1.0 | 1.2 | 1.0 | 1.0 | 0.8 | 0.9 | 1.0 | 1.1 |

### Software implementation, memory requirement and computational complexity

We implemented our algorithms in a software package, microbiomeGWAS, which is freely available at https://github.com/lsncibb/microbiomeGWAS. MicrobiomeGWAS requires three sets of files: a microbiome distance matrix file, a set of PLINK binary files for GWAS genotypes, and a set of covariates. MicrobiomeGWAS processes one SNP at a time and does not load all genotype data into memory; thus, it requires only memory for storing the distance matrix. Variance, skewness and kurtosis can be partitioned into two parts related with the microbiome distance matrix and the MAF of the SNP separately; thus, we can quickly calculate these quantities for a predefined grid of MAFs. The overall computational complexity is about *O*(*N*^2^*M*), where *N* is sample size and *M* is the number of SNPs. Figure 4 reports the computation time on a Linux server using a single core.

**Figure 4.**
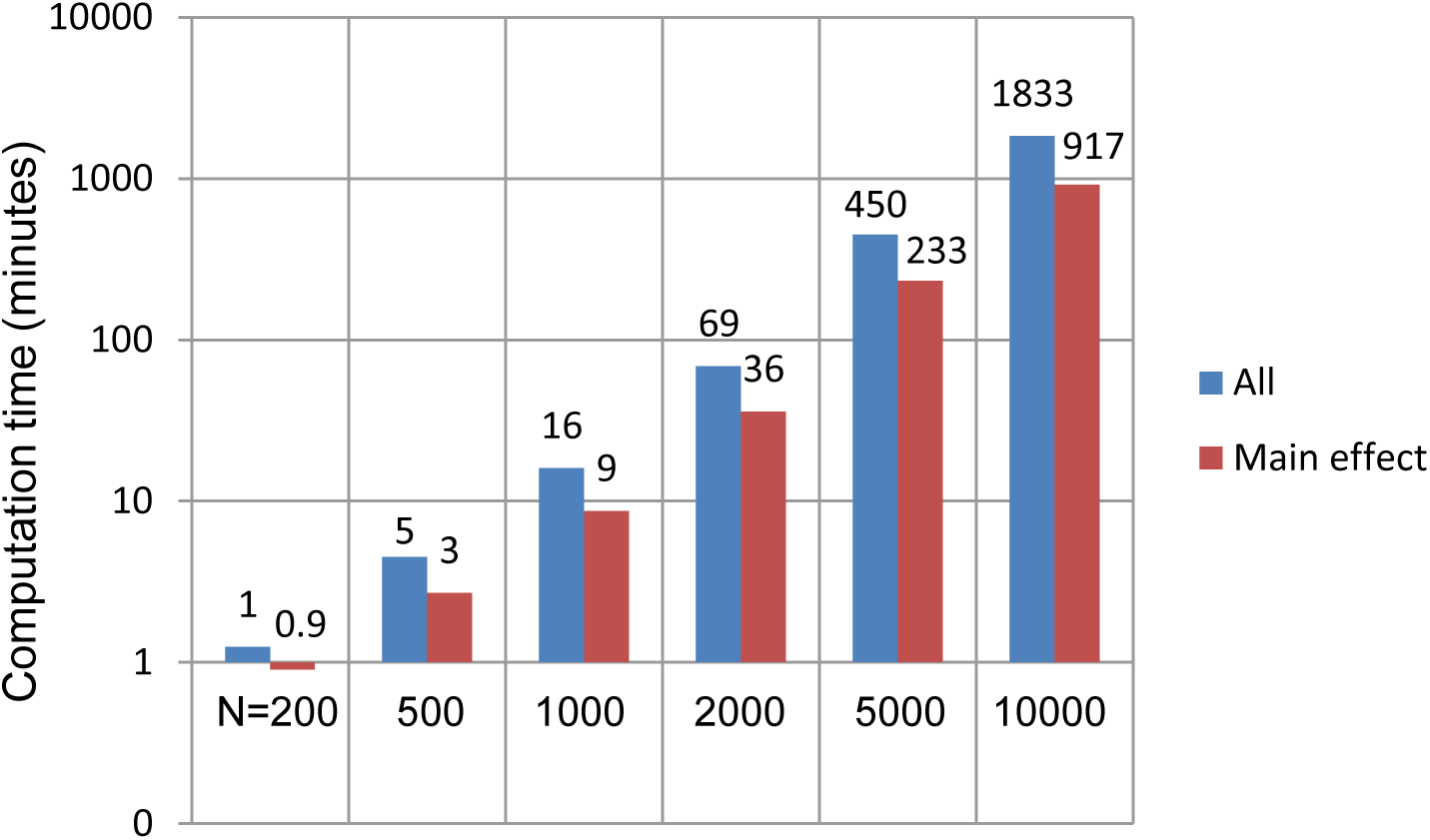
Computation time for a microbiome GWAS with 500,000 SNPs. “Main”: computation time for testing main effect only. “All”: computation time for testing main effect, interaction and the joint null hypothesis *H*_0_: *β_M_* = 0, *β_I_* = 0.

### GWAS of microbiome diversity in adjacent normal lung tissues

We applied our methods to a set of lung cancer patients of Italian ancestry in the EAGLE^28^ study. All subjects have germline genome-wide SNPs^29^ and 16S rRNA microbiome data (V3-V4 region, Illumina MiSeq, 300 paired-end) in histologically normal lung tissues from these patients. Here, the histologically normal lung tissues were 1~5 cm from the tumor tissue. We performed a series of quality control steps to filter out low quality sequence reads: average quality score <20 over 30bp windows, less than 60% similarity to the Greengenes^32^ reference or identified as chimera reads using UCHIME^33^. Sequence reads were then processed by QIIME^21^ to produce relative abundances (RA) of taxa, two alpha diversity metrics (observed number of species and Shannon’s index) and beta-diversity metrics (unweighted and weighted UniFrac distances) rarified to 1000 reads. We included 147 subjects with at least 1000 high quality sequence reads for genetic association analysis.

Out of the 147 subjects, 78 are current smokers, 8 are never smokers and 61 are former smokers. Because of the small number of never smokers, we merged never and former smokers as non-current smokers. All of the genetic association analyses were adjusted for sex, age, smoking status, and the top three PCA scores derived based on genome-wide SNPs. Here, the top three PCA scores were selected controlling population stratification because other PCA scores were unassociated with the distance matrices. We included 383,263 common SNPs with MAF ≥ 10% because rarer SNPs were expected to have no statistical power given the current sample size. We first performed GWAS analysis using PLINK^34^ to identify SNPs associated with taxa with average RA greater than 0.1% or two alpha-diversity metrics. We did not detect genome-wide significant associations with either main effects or gene by smoking interactions.

Next, we performed GWAS analysis using unweighted and weighted UniFrac distance matrices as a representation of eubacteria beta-diversity. The results for testing main effects are reported in **Figure 5**. Results for testing joint effects (main effect and SNP by smoking status interaction) are reported in **Figure S1**. Because of the small sample size, we observed large values of skewness and kurtosis with magnitude varying with the MAF of the SNPs (**Figure 5A**). The score statistics based on the weighted UniFrac distance matrix had a much larger skewness and kurtosis than did the unweighted UniFrac matrix. **Figures 5B and 5C** report the quantile-quantile (QQ) plot of the logarithm of the association P-values for the unweighted and weighted UniFrac distance matrices, respectively. For each distance matrix, we produced QQ plots for P-values based on the asymptotic approximation and for P-values adjusted for skewness and kurtosis. For both distance matrices, the QQ plots before adjustment strongly deviated from the expected uniform distribution. Our adjustment eliminated the deviation. In addition, consistent with the observation that the skewness and kurtosis were larger for the weighted UniFrac distance matrix, the QQ plot deviated more for the analysis based on the weighted UniFrac distance. Note that the skewness and kurtosis only affect the tail probabilities; thus, the inflation of the QQ plot is not reflected by the genomic control lambda value^35^ calculated as the median of P-values. In fact, lambda ≈ 1 for all four QQ plots.

**Figure 5.**
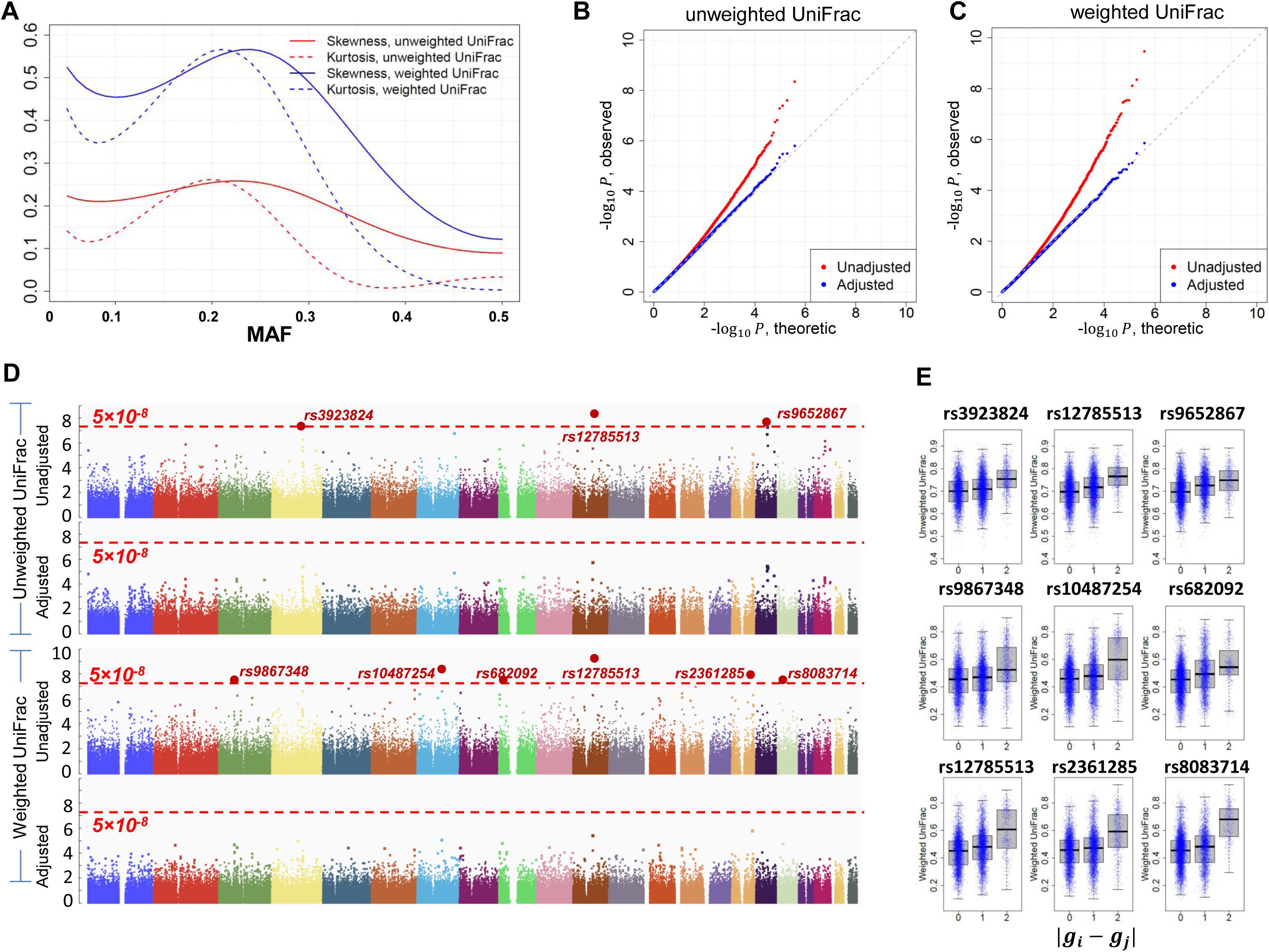
Results of analyzing the microbiome GWAS data of 147 adjacent normal lung tissues in the EAGLE study. (A) Skewness and kurtosis for the main effect test using the unweighted and the weighted UniFrac distance matrices. (B) Quantile-quantile (QQ) plot for association P-values using the unweighted UniFrac distance matrix. “Adjusted”: P-values were corrected for skewness and kurtosis. “Unadjusted”: P-values were approximated based on the asymptotic distribution *N*(0,l). (C) Quantile-quantile (QQ) plot for association P-values using the weighted UniFrac distance matrix. (D) Manhattan plots based on the unweighted or the weighted UniFrac distance matrices. (E) Box plots for the top nine loci in microbiome GWAS analysis. Subject pairs are classified into three groups according to the genetic distance |*g_i_* − *g_j_*| at the SNP. The y-coordinate is the microbiome distance.

Without correcting for skewness and kurtosis, we identified three and six loci achieving genome-wide significance (*P* < 5 × 10^−8^) for the unweighted and weighted UniFrac distance matrices, respectively (**Figure 5D**). After correcting for skewness and kurtosis, no locus remained genome-wide significant (**Figure 5D**), which was verified by 10^8^ permutations. Importantly, skewness and kurtosis had a dramatic effect on tail probabilities. Here, we use SNP rs12785513 as an example, which was identified as the top SNP in both analyses. In the unweighted UniFrac analysis, P= 4.4×10^−9^ without adjustment and P=1.6×10^−6^ after adjustment, a 364-fold inflation. The inflation was even larger for weighted UniFrac analysis because of larger skewness and kurtosis (**Figure 5A**). In fact, P=3.4×10^−10^ without adjustment and P=3.5×10^−6^ after adjustment, a 1000-fold inflation. Although these SNPs were not significant genome-wide, they were the top SNPs from the current study. Thus, we report box-plots for each of these nine SNPs (**Figure 5E**). As expected, in all box plots, microbiome distances tend to be larger in subject pairs with greater genetic distance at these SNPs. These associations remain to be replicated in studies with larger sample sizes.

Finally, we concentrated on the six common SNPs in four genomic regions reported to be associated with lung cancer risk in GWAS of European subjects: rs2036534 and rs1051730 at 15q25.1^36-39^ (*CHRNA5–CHRNA3–CHRNB4*), rs2736100 and rs401681 at locus 5p15.33^29,40^ (*TERT/CLPTM1L*), rs6489769^41^ at 12p13.3 (*RAD52*), and rs1333040 at 9p21.3^42^ (*CDKN2A/CDKN2B*). The SNPs at 15q25.1 and 5p15.33 have the largest effect sizes for lung cancer risk based on the meta-analysis from the Transdisciplinary Research in Cancer of the Lung (TRICL) consortium^42^: OR=1.32 for rs1051730, OR=1.26 for rs2036534, OR=1.13 for rs2736100, and OR=1.14 for rs401681. Rs3131379 at locus 6p21.33^40^ (*BAT3/MSH5*) was excluded because MAF=7.5%. No SNPs were significantly associated with taxa RAs or alpha-diversity metrics after correcting for multiple testing (data not shown). However, association analysis based on the UniFrac distance matrices provided evidence that these SNPs may be associated with the lung microbiota (**Table 2**). Importantly, for both unweighted and weighted UniFrac analyses, all six SNPs had *P*-value < 0.5. These SNPs were independent except that rs2036534 and rs1051730 at 15q25.1 were weakly correlated with R^2^=0.15. A test combining six *z*-scores (*Z_M_*) and adjusting for the weak correlation yielded overall P-values 0.0033 and 0.011 for the unweighted and the weighted UniFrac distance matrices, respectively. These results suggest that lung cancer risk SNPs were enriched for genetic association with the composition of the lung microbiome. The results for testing interactions and joint effects are reported in **Table S2**.

**Table 2:**
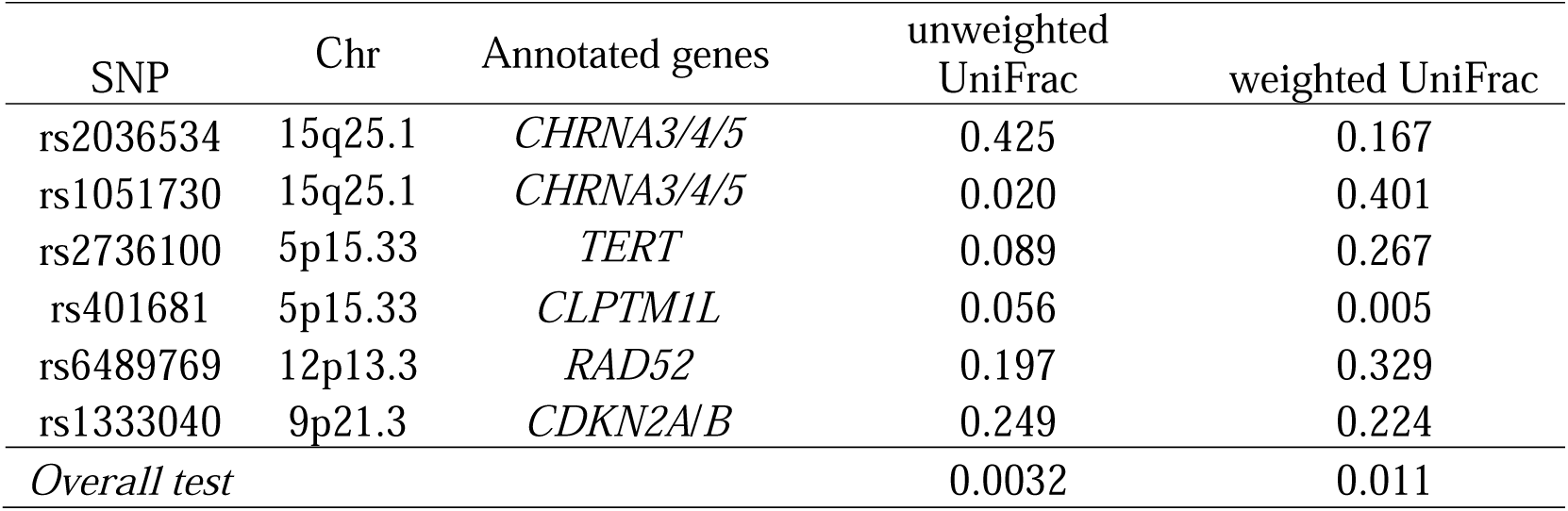
Association P-values between lung cancer risk SNPs and microbiome composition in the EAGLE data.

### Discussion

We developed a software package, microbiomeGWAS, for identifying host genetic variants associated with microbiome composition. MicrobiomeGWAS can test both main effect and SNP-environment interactions. Importantly, we found that the score statistics had positive skewness and kurtosis and that the tail probabilities evaluated based on asymptotic approximations were very liberal. We addressed this problem by explicitly adjusting for skewness and kurtosis. MicrobiomeGWAS runs very efficiently and takes only 36 minutes for testing main effects and 69 minutes for testing joint effects in a GWAS with 2000 subjects and 500,000 markers. Other statistical methods exist for testing the association of microbiome distance matrices. PERMANOVA^43^ is an extension of multivariate analysis of variance to a matrix of pairwise distances and relies on permutations to evaluate significance. MiRKAT^44^, a recently proposed method based on kernel regression, takes hours for evaluating one association for 2000 subjects. Neither is computationally feasible for analyzing a large-scale GWAS of microbiome.

Interactions of host genetic susceptibility with the microbiome have been postulated for many conditions, including inflammatory bowel diseases^45,46^, autoimmune and rheumatic diseases^47-50^, diabetes^51^, and cancer especially of the colon^52^. All models of these host-microbiome interactions also note the critical role of environmental factors including diet, smoking, drugs, and antibiotics and other medications^53^.

Although based on a very small initial sample set, the suggestive associations that we found between the six known lung cancer risk SNPs and the microbiome of adjacent normal lung tissue samples, including effects of cigarette smoking, provide preliminary evidence that our microbiomeGWAS method is likely to be a useful tool for generating data that will unravel host-microbiome interactions with high confidence.

We are working on two extensions for microbiomeGWAS: (1) jointly testing additive and dominant effects and (2) testing genetic associations using many microbiome distance matrices. We have assumed an additive effect model (**Figure 1**); however, several top SNPs in the EAGLE data suggest a dominant effect (e.g. rs8083714 in **Figure 5E**). Thus, a statistic for jointly testing the additive and dominant effects might be powerful for this scenario. The second extension is motivated by the fact that that the power to detect associations depends heavily on the choice of distance matrix. The recently developed generalized UniFrac^26^ (gUniFrac) defines a series of distance matrices to reflect different emphasis of using taxa relative abundance information. gUniFrac has been shown to have a robust power for association studies^26^. Extending microbiomeGWAS to gUniFrac, however, requires solving two problems. First, the computational complexity is proportional to the number of distance matrices analyzed for associations, which can be addressed by implementing the algorithms using multithreading technology. Second, we need to derive accurate analytic approximations to the association *P*-values by correcting for the multiple testing introduced by many distance matrices. MiRKAT^44^ has an option for using gUniFrac; however, intensive permutations are required to evaluate P-values.

In summary, GWAS of the microbiome of each body site has a potential to understand microbiome variation, to elucidate biological mechanisms of genetic associations, to improve the power of identifying novel disease-associated genetic variants, and to improve the performance of genetic risk prediction. We expect our methods and software to be useful for large-scale GWAS of human microbiome.

## Appendices

## Appendix A: *Var*(*G_ij_*), *Cov*(*G_ij_,G_ifc_*), *Var*(Δ*_ij_*) and *Cov*(Δ*_ij_*, Δ*_ik_*).

We first calculate *E*(*G_ij_*), *Var*(*G_ij_*) and *Cov*(*G_ij_*, *G_ik_*). Let *p_t_* = *P*(*g_j_* = *t*) with *p*_0_*, p*_1_*, p*_2_ ≥ 0 and *p*_0_ *+ p*_1_ *+ p*_2_ = 1. We can also assume the Hardy–Weinberg equilibrium and characterize the probabilities as the allele frequency: *p*_0_ = (1 − *f*)^2^, *p*_1_ = 2*f*(1 − *f*) and *p*_2_ = *f*^2^. Some algebra leads to

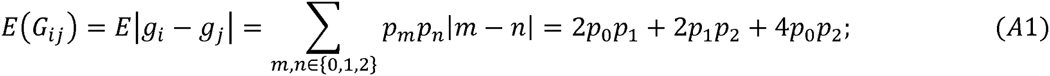

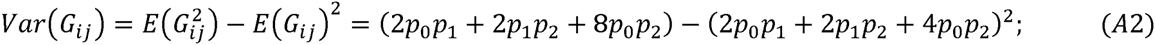

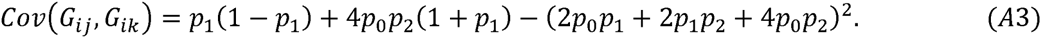

Now consider Δ*_ij_*·= |*g_i_E_i_* – *g_j_E_j_*|. When *E_i_* is binary, *g_i_E_i_* = 0,1 or 2. Let 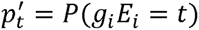. Then, *E*(Δ*_ij_*), *Var*(Δ*_ij_*) and *Cov*(Δ*_ij_*, Δ*_ik_*) can be calculated similarly using (A1), (A2) and (A3).ρ

## Appendix B: Calculating *ρ = Cor*_0_(*Z_M_*,*Z_I_\****D***′)

Let 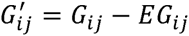 and 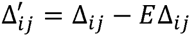. We first calculate the covariance under *H*_0_:

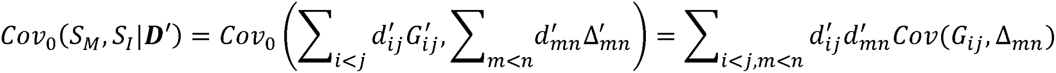

When (*i, j, m, n*) are distinct, *Cov*(*G_ij_*, Δ*_mn_*) = 0. Some algebra leads to

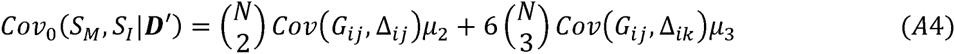

with *μ*_2_ and *μ*_3_ specified in (2) and (3). Combining (2), (5) and (A4), we have

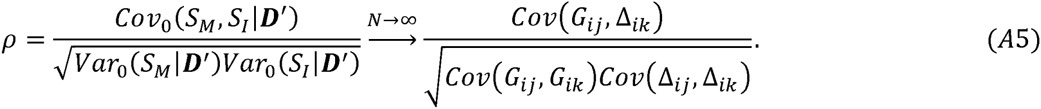

(A5) suggests that the correlation is asymptotically independent of the microbiome distance matrix. In real data analyses, we found that (A5) was very accurate when sample size *N* ≥ 50. The details of calculating and are provided in **Supplemental Data**.

## Appendix C: A statistic for jointly testing *H_0_·. β_M_ = β_I_* = 0 vs *H*_1_:*β_M_* > 0 or *β_I_* > 0

Denote *Z* = (*Z_M_*, *Z_I_*)*^T^*. Under *H*_0_, ***Ζ****~Ν*(***0, Σ***) with 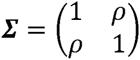. Let *ξ_Μ_* = *E*_1_*Z_M_ ≥* 0 and *ξ_Ι_ = E*_1_*Z_I_* ≥ 0 be the non-centrality parameter of the two score statistics. Apparently the original testing problem is equivalent for testing *H*_0_*: ξ_Μ_* = *ξ_Ι_* = 0 vs *H*_1_*. ξ_Μ_* > 0 or *ξ_Ι_* > 0. Given the observed values (*Z_M_,Z_I_*), the likelihood ratio statistic is simplified as

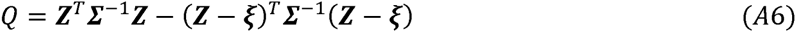

where 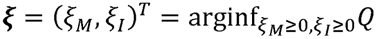 (**Figure S2A**).

To simplify the optimization problem in (A6), we perform a linear transformation: 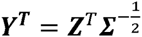 and 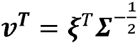, where

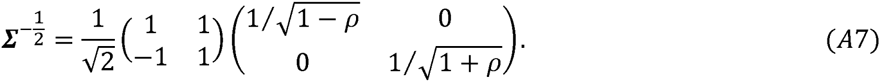

Under this transformation, *Q = Y^T^Y* − (*Y* − *v*)*^T^*(*Y* − *v*) and can be interpreted as the difference of the square of two distances (**Figure S2B**). The original parameter space {(*ξ_Μ_, ξ_I_*)*·ξ_Μ_* ≥ 0, *ξ_I_* ≥ 0} is now transformed to {(*ν*_1_, *v*_2_): *v*_2_ ≥ *θν*_1,_*v*_2_ *≥ −θv*_1_} with 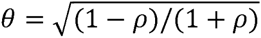. Thus, the new parameter space is bounded by two lines represented by *v*_2_ ≥ *θv*_1_ and *v*_2_ ≥ −*θν*_1_. We partition the 2D plane into four parts (see **Figure S2B**), identify 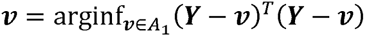 and calculate *Q*:

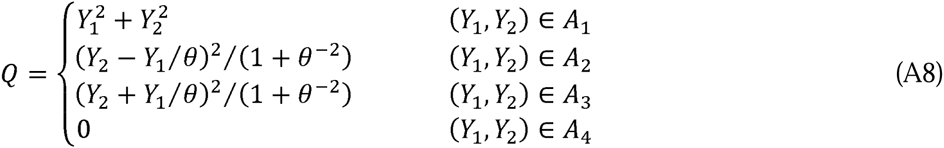

We now perform an inverse transformation using matrix

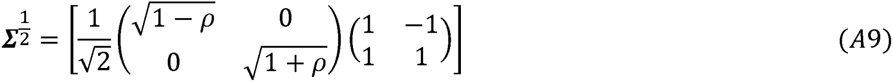

to return to the original parameter space. The four areas {*Α*_1_*, A*_2_*, A*_3_*, A*_4_} under the original space are in **Figure 2** and **Figure S2C**.

Tedious calculations show that with (*Y_2_ + Υ_1_/θ*)^2^*/*(1 *+ θ*^−2^) = (*w*_2_*Z_M_ + w*_1_*Z_I_*)^2^ with *w*_1_ = (*θ −* l/*θ*)/2 and *w*_2_ = (*θ +* l/*θ*)/2. Similarly, (*Y_2_ − Υ_1_/θ*)^2^*/*(1 *+ θ^−^*^2^) = (*w*_1_*Z_M_ + w*_2_*Z_I_*)^2^ *>.* This proves (7). In addition, *w_1_Z_M_ + w_2_Z_I_* ≥ 0 and 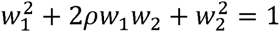 thus, *P*{(*w*_1_*Z_M_ + w*_2_*Z_I_*)^2^ *> b^2^*} *= P*{*w_1_Z_M_ + w_2_Z_I_ > b*} *= P*{*N*(0,1) > *b*}. This proves (8). The probabilities in (8) could also be calculated from **Figure S2B**: *q*_1_ = 1/2 − (arctan *θ*)*/π,q*_2_ *= q*_3_ = 1/4.

## Appendix D: Calculating skewness and kurtosis under *H*_0_

By definition, 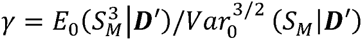 and 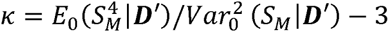. We first calculate 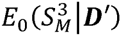. Let 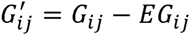. We have

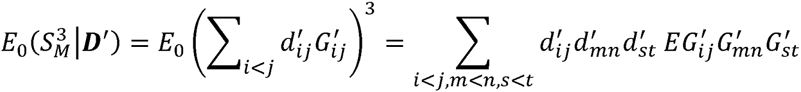

**Figure S3A** lists all combinations of (*i, j, m, n, s, t*) with 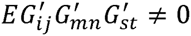, which leads to

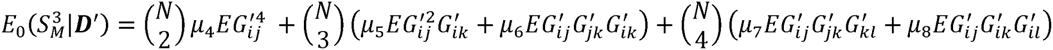

where (*μ*_4_, *μ*_5_, *μ*_6_, *μ*_7_, *μ*_8_) are provided in **Supplemental Data**. Similarly,

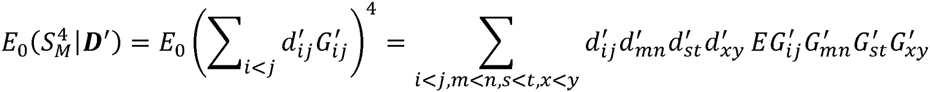

**Figure S3B** lists 15 combinations of (*i,j,m,n,s,t,x,y*) with 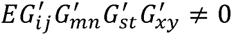. Thus,

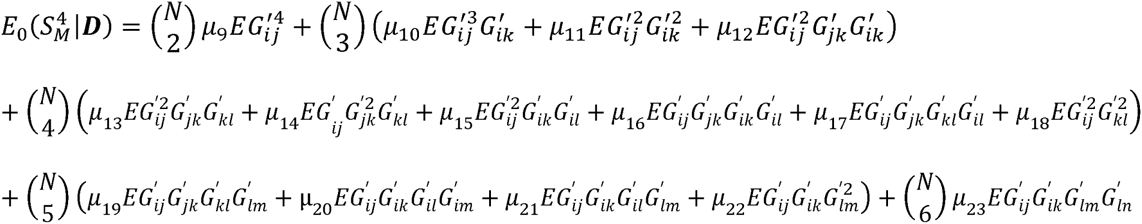

The constants (*μ*_9_, …,*μ*_23_) are dependent on ***D*** and are provided in **Supplemental Data**.

Note that *Var*_0_(*S_M_\****D****′*)*~O*(*N^3^*), 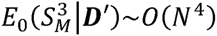, thus 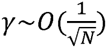. Similarly, we can prove 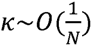.

## Appendix E: Improve *P*-value approximations by adjusting for skewness and kurtosis

We assume that *E*_0_*Z* = 0, *Var*_0_*Z* = 1, γ = *E*_0_*Z*^3^ and *k* = *E*_0_*Z*^4^− 3 under the original probability measure *P*_0_. The tail probability *P*_0_(*Z > b*) for a large value of *b* is sensitive to the non-normality of *Z*, characterized by *γ* and *κ.* We define a new probability measure by embedding to the exponential probability density

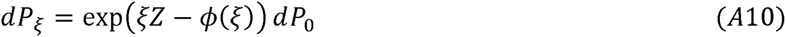

where *ϕ*(*ξ*) = log *E*_0_ exp(*ξZ*) is the log moment generating function Note that γ = *ϕ*′″(0) and *k* = *ϕ*′″(0). Because *E*_0_ (*Z*) = 0 and *Var*_0_ (*Z*) = 1, the Taylor expansion leads to *ϕ*(*ξ*) ≈ *ξ*^2^/2 + γ*ξ*^3^/6 + *kξ*^4^/24. Under *P_ξ_*, we have

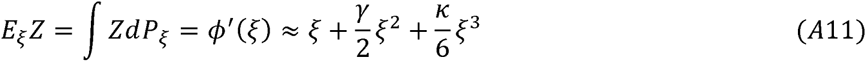

and

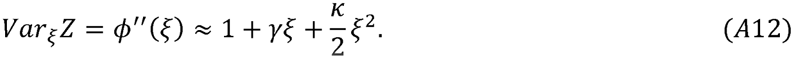

We choose *ξ* such that *E_ξ_Z ≈ b* by numerically solving an equation

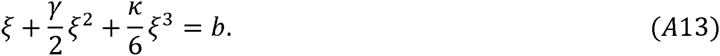

Under the probability measure *Ρ_ξ_, Z~N*(*b, σ^2^*) approximately with *σ^2^* = 1 + *γξ + kξ^2^*/*2* in (A12).

By the likelihood ratio identity and (A10), we have

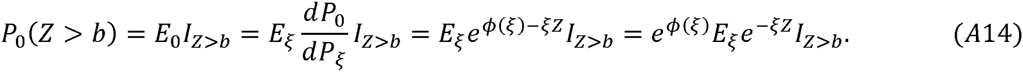

Note that *e^−ξz^* decays very fast when *Z* increases. Thus, the integral *Ε_ξ_e^−ξz^I_z>b_* does not heavily depend on the tail distribution of *Z*. Assuming *Z ~ N*(*b, σ*^2^) under *P_ξ_*, we can verify that

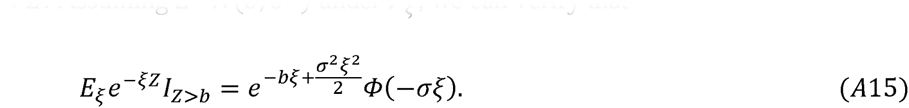

Combining (A14) and (A15) gives 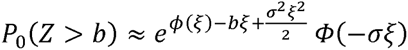, which is further approximated as 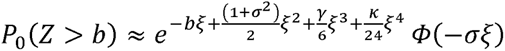, because *ϕ*(*ξ*) *≈ ξ*^2^/2 + γ*ξ*^3^/6 + *κξ*^4^/24 based on the Taylor expansion. This proves (9).

If we correct skewness but assume kurtosis *k* = 0, then *ϕ*(*ξ*) *≈ ξ*^2^/2 + γ*ξ*^3^/6. We recalculate *ξ* by setting *κ* = 0 in (A13) to derive 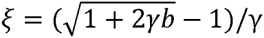. This proves (10).

## Supplemental Data

Supplemental Data include 2 tables and 3 figures can be found with this article online at XXX.

## Acknowledgements

This study utilized the high-performance computational capabilities of the Biowulf Linux cluster at the National Institutes of Health, Bethesda, MD. (http://biowulf.nih.gov). The project was supported by the NIH Intramural Research program.

## Web Resources

The URLs for data provide herein are as follows:

American Gut Project: https://www.microbio.me/americangut/

PLINK: http://pngu.mgh.harvard.edu/~purcell/plink/

QIIME: http://qiime.org/

EAGLE study: http://eagle.cancer.gov/

## References

1. Turnbaugh, P.J. et al. A core gut microbiome in obese and lean twins. Nature 457, 480–4 (2009).

2. Morgan, X.C. et al. Dysfunction of the intestinal microbiome in inflammatory bowel disease and treatment. Genome Biol 13, R79 (2012).

3. Ahn, J. et al. Human gut microbiome and risk for colorectal cancer. J Natl Cancer Inst 105, 1907–11 (2013).

4. Goedert, J.J., Jones, G., Hua, X., Xu, X., Yu, G., Flores, R., Falk, R. T., Gail, M. H., Shi, J., Ravel, J. and Feigelson, S. H. Investigation of the Association Between the Fecal Microbiota and Breast Cancer in Postmenopausal Women: a Population-Based Case-Control Pilot Study. J Natl Cancer Inst. 1;107(8)(2015).

5. Lax, S. et al. Longitudinal analysis of microbial interaction between humans and the indoor environment. Science 345, 1048–1052 (2014).

6. Wu, G.D. et al. Linking long-term dietary patterns with gut microbial enterotypes. Science 334, 105–8 (2011).

7. Tong, M. et al. Reprograming of gut microbiome energy metabolism by the FUT2 Crohn’s disease risk polymorphism. ISME J 8, 2193–206 (2014).

8. Knights, D. et al. Complex host genetics influence the microbiome in inflammatory bowel disease. Genome Med 6, 107 (2014).

9. McKnite, A.M. et al. Murine Gut Microbiota Is Defined by Host Genetics and Modulates Variation of Metabolic Traits. Plos One 7(2012).

10. Benson, A.K. et al. Individuality in gut microbiota composition is a complex polygenic trait shaped by multiple environmental and host genetic factors. Proc Natl Acad Sci U S A 107, 18933–8 (2010).

11. Goodrich, J.K. et al. Human genetics shape the gut microbiome. Cell 159, 789–99 (2014).

12. Davenport, E.R. et al. Genome-Wide Association Studies of the Human Gut Microbiota. PLoS One 10, e0140301 (2015).

13. Consortium, G.T. Human genomics. The Genotype-Tissue Expression (GTEx) pilot analysis: multitissue gene regulation in humans. Science 348, 648–60 (2015).

14. Battle, A. et al. Characterizing the genetic basis of transcriptome diversity through RNA-sequencing of 922 individuals. Genome Research 24, 14–24 (2014).

15. Bell, J.T. et al. DNA methylation patterns associate with genetic and gene expression variation in HapMap cell lines (vol 12, pg R10, 2011). Genome Biology 12(2011).

16. Shi, J. et al. Characterizing the genetic basis of methylome diversity in histologically normal human lung tissue. Nat Commun 5, 3365 (2014).

17. McVicker, G. et al. Identification of Genetic Variants That Affect Histone Modifications in Human Cells. Science 342, 747–749 (2013).

18. Kilpinen, H. et al. Coordinated Effects of Sequence Variation on DNA Binding, Chromatin Structure, and Transcription. Science 342, 744–747 (2013).

19. Suhre, K. et al. A genome-wide association study of metabolic traits in human urine. Nat Genet 43, 565–9 (2011).

20. Sabatti, C. et al. Genome-wide association analysis of metabolic traits in a birth cohort from a founder population. Nat Genet 41, 35–46 (2009).

21. Caporaso, J.G. et al. QIIME allows analysis of high-throughput community sequencing data. Nat Methods 7, 335–6 (2010).

22. Bray, J.R.a.C., J. T. An ordination of upland forest communities of southern Wisconsin. Ecological Monographs 27:325–349(1957).

23. Lozupone, C. & Knight, R. UniFrac: a new phylogenetic method for comparing microbial communities. Appl Environ Microbiol 71, 8228–35 (2005).

24. Lozupone, C.A., Hamady, M., Kelley, S.T. & Knight, R. Quantitative and qualitative beta diversity measures lead to different insights into factors that structure microbial communities. Appl Environ Microbiol 73, 1576–85 (2007).

25. Lozupone, C., Hamady, M. & Knight, R. UniFrac-an online tool for comparing microbial community diversity in a phylogenetic context. BMC Bioinformatics 7, 371 (2006).

26. Chen, J. et al. Associating microbiome composition with environmental covariates using generalized UniFrac distances. Bioinformatics 28, 2106–13 (2012).

27. Gevers, D. et al. The Human Microbiome Project: a community resource for the healthy human microbiome. PLoS Biol 10, e1001377 (2012).

28. Landi, M.T. et al. Environment And Genetics in Lung cancer Etiology (EAGLE) study: an integrative population-based case-control study of lung cancer. BMC Public Health 8, 203 (2008).

29. Landi, M.T. et al. A genome-wide association study of lung cancer identifies a region of chromosome 5p15 associated with risk for adenocarcinoma. Am J Hum Genet 85, 679–91 (2009).

30. Tu, I.P. & Siegmund, D. The maximum of a function of a Markov chain and application to linkage analysis. Advances in Applied Probability 31, 510–531 (1999).

31. Siegmund, D. Sequential Analysis: Tests and Confidence Intervals. New York: Springer. (1985).

32. DeSantis, T.Z., Dubosarskiy, I., Murray, S.R. & Andersen, G.L. Comprehensive aligned sequence construction for automated design of effective probes (CASCADE-P) using 16S rDNA. Bioinformatics 19, 1461–8 (2003).

33. Edgar, R.C., Haas, B.J., Clemente, J.C., Quince, C. & Knight, R. UCHIME improves sensitivity and speed of chimera detection. Bioinformatics 27, 2194–200 (2011).

34. Purcell, S. et al. PLINK: a tool set for whole-genome association and population-based linkage analyses. Am J Hum Genet 81, 559–75 (2007).

35. Devlin, B. & Roeder, K. Genomic control for association studies. Biometrics 55, 997–1004 (1999).

36. Hung, R.J. et al. A susceptibility locus for lung cancer maps to nicotinic acetylcholine receptor subunit genes on 15q25. Nature 452, 633–7 (2008).

37. McKay, J.D. et al. Lung cancer susceptibility locus at 5p15.33. Nat Genet 40, 1404–6(2008).

38. Thorgeirsson, T.E. et al. A variant associated with nicotine dependence, lung cancer and peripheral arterial disease. Nature 452, 638–42 (2008).

39. Amos, C.I. et al. Genome-wide association scan of tag SNPs identifies a susceptibility locus for lung cancer at 15q25.1. Nat Genet 40, 616–22 (2008).

40. Wang, Y. et al. Common 5p15.33 and 6p21.33 variants influence lung cancer risk. Nat Genet 40, 1407–9 (2008).

41. Shi, J. et al. Inherited variation at chromosome 12p13.33, including RAD52, influences the risk of squamous cell lung carcinoma. Cancer Discov 2, 131–9 (2012).

42. Timofeeva, M.N. et al. Influence of common genetic variation on lung cancer risk: meta-analysis of 14 900 cases and 29 485 controls. Hum Mol Genet 21, 4980–95 (2012).

43. Anderson, M.J. A new method for non-parametric multivariate analysis of variance. Austral Ecology 26:, 32–46. (2001).

44. Zhao, N. et al. Testing in Microbiome-Profiling Studies with MiRKAT, the Microbiome Regression-Based Kernel Association Test. Am J Hum Genet 96, 797–807 (2015).

45. Leone, V.A., Cham, C.M. & Chang, E.B. Diet, gut microbes, and genetics in immune function: can we leverage our current knowledge to achieve better outcomes in inflammatory bowel diseases? Current Opinion in Immunology 31, 16–23 (2014).

46. Huang, H., Vangay, P., McKinlay, C.E. & Knights, D. Multi-omics analysis of inflammatory bowel disease. Immunol Lett 162, 62–8 (2014).

47. Troncone, R. & Discepolo, V. Celiac disease and autoimmunity. J Pediatr Gastroenterol Nutr 59 Suppl 1, S9-S11 (2014).

48. Yeoh, N., Burton, J.P., Suppiah, P., Reid, G. & Stebbings, S. The role of the microbiome in rheumatic diseases. Curr Rheumatol Rep 15, 314 (2013).

49. Sparks, J.A. & Costenbader, K.H. Genetics, environment, and gene-environment interactions in the development of systemic rheumatic diseases. Rheum Dis Clin North Am 40, 637–57 (2014).

50. Smith, J.A. Update on ankylosing spondylitis: current concepts in pathogenesis. Curr Allergy Asthma Rep 15, 489 (2015).

51. Nielsen, D.S., Krych, L., Buschard, K., Hansen, C.H. & Hansen, A.K. Beyond genetics. Influence of dietary factors and gut microbiota on type 1 diabetes. FEBS Lett 588, 4234–43 (2014).

52. Birt, D.F. & Phillips, G.J. Diet, genes, and microbes: complexities of colon cancer prevention. Toxicol Pathol 42, 182–8 (2014).

53. Marietta, E., Rishi, A. & Taneja, V. Immunogenetic control of the intestinal microbiota. Immunology 145, 313–22 (2015).

